# Mapping the Landscape of Allele-Specific Expression In Porcine Genomes

**DOI:** 10.1101/2025.02.19.639037

**Authors:** Wen-ye Yao, Marta Gòdia, Lingzhao Fang, Martien A. M. Groenen, Lijing Bai, Kui Li, Ole Madsen

## Abstract

Allele-specific expression (ASE) is the imbalanced expression of two alleles of the same locus. It is quite pervasive among mammals and is associated with healthy and economically relevant traits. ASE is often used to support the identification of variants related to gene expression (cis-eQTLs). Thus, profiling ASE represents a significant step in elucidating the mechanism underlying gene expression regulation. In this study, we developed an ASE pipeline using public available RNA-seq data and open-source software. Using this pipeline, we were able to profile pervasive allelic imbalance across 42 tissues and 34 breeds from the Farm-GTEX-pig consortium at both SNP and gene levels without the need for parental genotype or whole genome sequence data. ASE was widely, but not evenly, spread across the genome. We also observed considerable variation in ASE profiles among various tissues, in which the ASE fraction ranged from 1.3% to 54.1%. ASE tends to be highly tissue-specific, and the overlap across tissues is limited. The functional analysis of tissue-specific ASE sites indicates that they are involved in the critical maintenance of these tissues. Our ASE pipeline can be readily applied to other RNA-seq data sets for livestock, thereby significantly expanding its potential utility. The wealth of available ASE resources provides a solid foundation for identifying regulatory elements within the genome that drive complex traits in livestock, making our pipeline and results valuable resources for researchers in this field.

## Background

Diploid organisms carry two alleles at each locus. For a long time, it was thought that both alleles were equally expressed and hereby equally contributing to an individual’s phenotype. Most of the time, the expression of two alleles is indeed equal, but for some genes, the expression of the two alleles is unequal, a phenomenon called allelic imbalance or allele-specific expression (ASE). ASE appears to be a relatively common phenomenon throughout mammalian genomes [1, 2]. The most well-known ASE phenomena include Chromosome X-inactivation [3], in which the inactivated chromosome is suppressed as a result of epigenetic changes in the chromosome structure, and imprinted genes [4], where epigenetic signatures govern the parent-dependent expression of the imprinted genes [5]. Multiple genetic and epigenetic mechanisms can lead to ASE, while most ASE is caused by genetic variation [6]. These genetic variants can impact the binding affinity of transcription factors [7], chromosomal states [8, 9], and mRNA splicing [10], leading to ASE. ASE can be identified and quantified by RNA sequencing, typically overlapping uniquely mapped RNA-sequence reads to heterozygous sites and counting expression level of each allele. Genotype data can be retrieved from DNA sequences, genotype arrays, and by imputation based on SNPs called from RNA-sequencing data.

The prevalence and pattern of ASE have been widely studied across tissues in human genomic analyses. Data from the human Lymphoblastoid cell line (LCL) by the Geuvadis consortium [7] suggested that 6.5% of the LCL genes show ASE. The prevalence of ASE across the human GTEx samples showed that the distribution of significant ASE fractions ranged from 1.7 to 3.7% across tissues [11]. This research indicated that brain regions appeared to have the lowest ASE fraction (∼ 2%), while the blood showed the highest ASE proportion (∼3.6%). A later human GTEx data release (GTEx v8), comprised of a comprehensive ASE resource derived from 15,253 samples from 54 tissues, showed that 53% of protein-coding genes showed strong ASE in at least 50 individuals [1]. Another analysis explored the changes in ASE over time from the same individuals at ages 70 and 80 [12], in which a 2.6% increase in ASE with ages was observed. Many genes that showed changes in ASE over time were associated with the immune response, suggesting that ASE could be involved in aging.

A relatively diverse ASE fraction was found among tissues of different mammals, such as cattle, sheep, mice, and pigs. Around 20% of mouse transcripts showed ASE in tail-tip fibroblasts [13]. Several studies have investigated the prevalence of ASE in cattle [2, 14–17]. These studies generally indicated that ASE is widely spread throughout cattle genomes across different sub-species while varying wildly among various tissues, ranging from 10% in the thymus to 65% in the lung [2]. In crossbred blackface sheep, the ASE discovery rate comprised, on average, 5.8% of the heterozygous sites in each individual [18]. However, the shared ASE profile across the four sheep tissues and two cell types was limited, suggesting ASE to be highly tissue-specific. Pigs (*Sus scrofa*) are key agricultural animals serving as the primary meat resource and biomedical model. DNA and RNA sequencing data of pigs have expanded rapidly in scale and volume during the past two decades. Around 10.6% of heterozygous SNPs (11,300 SNPs) exhibited ASE in the prenatal skeletal muscle of Tongcheng pigs, with 4 of these ASE variants reported to be associated with average daily gain in a GWAS [19]. ASE analyses in peripheral blood identified 2,286 ASE sites from large white pigs, with functional enrichment related to RNA processing and immune function [20]. Furthermore, around 42% of all genes showed ASE in the brain [21]. Although these ASE analyses revealed the prevalence and importance of ASE in pigs, a substantial cohort study to explore ASE is still lacking despite thousands of RNA-seq data being available online. A more comprehensive analysis is essential for deeply investigating the regulatory mechanism behind critical economic traits in pigs.

This study provides ASE profiling from a comprehensive pig RNA-seq dataset containing 6,022 samples spanning 42 tissues (including 3 cell lines) and 34 breeds from the Farm-GTEx-pig project [22]. From this ASE profile, we provide insight into the distribution of ASE in the pig genome across tissues and breeds and generate tissue-shared and specific ASE profiles. This extensive ASE atlas data set provides a valuable foundation for further analyses to identify the molecular regulatory variants underlying complex traits in pigs.

## Results

### Genotype data comparison and mapping bias filtration

Variants from whole genome sequences are considered the gold standard for ASE detection. In our study, the genotype data were based on RNA called heterozygous SNPs imputed to the population level from the Farm-GTEx-pig consortium (see Methods). Such data has been proven successful for ASE detection despite its higher rate of false heterozygotes compared to WGS [23, 24]. To test the reliability of the imputed SNPs in our project, we compared the SNPs obtained from WGS and their corresponding RNA-imputed genotype data among 17 samples (derived from 7 tissues and 3 breeds; Additional file 2: Table S1). After filtering with the standard quality control criteria, the detected variants from WGS ranged from 7,153,561 to 8,504,747, and the variants from RNA-imputed genotype data ranged from 1,151,736 to 1,328,882 for each sample (Additional file 2: Table S1). On average, 86.97% of the RNA-imputed SNPs were also detected by WGS (Additional file 2: Table S1), and if two outliers (see below) were excluded, this similarity increased to 92.23%. We observed two outliers in the accuracy of the RNA-imputed genotype data, namely for the liver sample in Pietrain and the muscle sample in Duroc. These two samples also displayed deviating numbers of RNA-imputed SNPs (Additional file 2: Table S2). From the Farm-GTEx consortium database, it appears that these samples were wrongly assigned to a breed and, therefore, imputed from an incorrect breed. This shows the importance of correct population assignment for genotype imputation. Overall, most samples were correctly assigned to the breed and showed high precision in the RNA-imputed genotype data.

Reference allele mapping bias remains a considerable issue in ASE analyses. We compared the effect of the mapping bias correction using the two different genotype resources. The effect of removing the reference mapping bias was assessed by reference ratio. The reference ratio decreased to around 0.5 using both genotype data after applying the filtering criteria described in Methods (Additional file 1: Fig S2A). We then measured the effect of ASE detection using these two genotype data. With more SNP sites in the WGS genotype dataset, there are more ASE sites detected by the WGS genotype data compared to RNA-imputed genotype data (on average 1,379 .vs. 803, Additional file 2:Table S2 & Additional file 1: Fig S2B). Meanwhile, the muscle sample in Duroc and the liver sample in Pietrain exhibited a lower ASE trend than other muscle and liver samples. This discrepancy may be explained by the inconsistencies in SNPs between the WGS and RNA-imputed genotype data for these two samples. Considering that approximately 15% of RNA-imputed SNPs were not validated by WGS SNPs, it is reasonable to assume that some noise is present in the RNA-imputed ASE sites. However, we observed similar ASE trends among breeds and tissues for the WGS and RNA-imputed genotype data. This suggests that any potential bias from the RNA-imputed genotyping is consistent across tissues and breeds, enabling the use of RNA-imputed genotype data for the subsequent analysis. After applying the basic QC and filtering criteria (Method & Additional file 1: Fig S1), a total of 6,022 samples were retained for the subsequent analyses, encompassing 39 tissues and 3 cell lines from 34 breeds (Additional file 2: Table S2). For the tissues, the median sample size was 78.5, ranging from 2 (cerebellum) to 1,416 (muscle) (Additional file 1: Fig S2C). Two main breed categories were included, namely cross and pure breeds. For the pure breeds, there were a total of 1,845 samples, including the most common commercial breeds, such as Yorkshire and Duroc, and different local breeds, such as Wuzhishan, Meishan, and Tibetan breeds (Additional file 1: FigS2D). Most samples were from crossbreeds (N=4,177), which contained a variety of combinations of different breeds (Additional file 1: FigS2E & Additional file 2: TableS2). The reference ratio across all samples is close to 0.5 (Additional file 1: FigS2C, D, E), with a slight bias towards the reference allele. Thus, despite applying various filtering steps, it should be noted that the allelic mapping bias cannot be entirely eliminated.

### Genome-wide ASE profiles

For the SNP level ASE analysis, a mean of 11,316 tested sites (ranging from 268 to 92,052 sites) was obtained from the RNA-imputed SNPs for ASE testing, accounting for 1.71% of the 660,261 RNA-imputed SNPs located in gene regions. On average, 850 sites showed significant ASE (Fig1B), accounting for 7.51% of the total tested sites. The ASE sites per sample ranged from 10 to 46,825, with the corresponding ASE fraction ranging from 0.23% to 77.2% (Additional file 3: Table S3). A total of 309,891 ASE sites were detected by at least one sample, with 104,078 ASE sites identified only in one sample (Additional file 1: Fig S3A).

The ASE sites were broadly but unevenly distributed across autosomal chromosomes (Fig 1C) and generally consistent with the gene distribution in the genome (Adjusted R^2^=0.4658, *p-*value ≤ 0.05, Additional file 1: Fig S3B). The number of ASE sites per 1 Mb segment was, on average, 136. For 22,9% of the segments (521 of 2,272 total segments), no ASE sites were detected. The majority of these segments (485 of 521) had no tested sites either and were primarily located in intergenic or non-expressed regions. For the rest of the segments with ASE sites (1,751 segments), there were 7 segments with a high density of ASE sites (ASE site count ≥ 1,000), while 439 segments had a very low density of ASE sites (ASE site count ≤ 42). Thirty-seven segments showed a 100% ASE ratio, with all tested sites being significant, but the number of ASE sites in these segments was low, ranging from 1 to 14. The ASE was not only unevenly distributed within chromosomes but also showed unequal enrichment across different chromosomes (Fig 1D). The highest ASE site density was on chromosome 12 after normalization for chromosome length, with a density of 366 sites per 1 Mb. The lowest density was on chromosome 16, with 77 sites per 1 Mb.

**Fig. 1.**
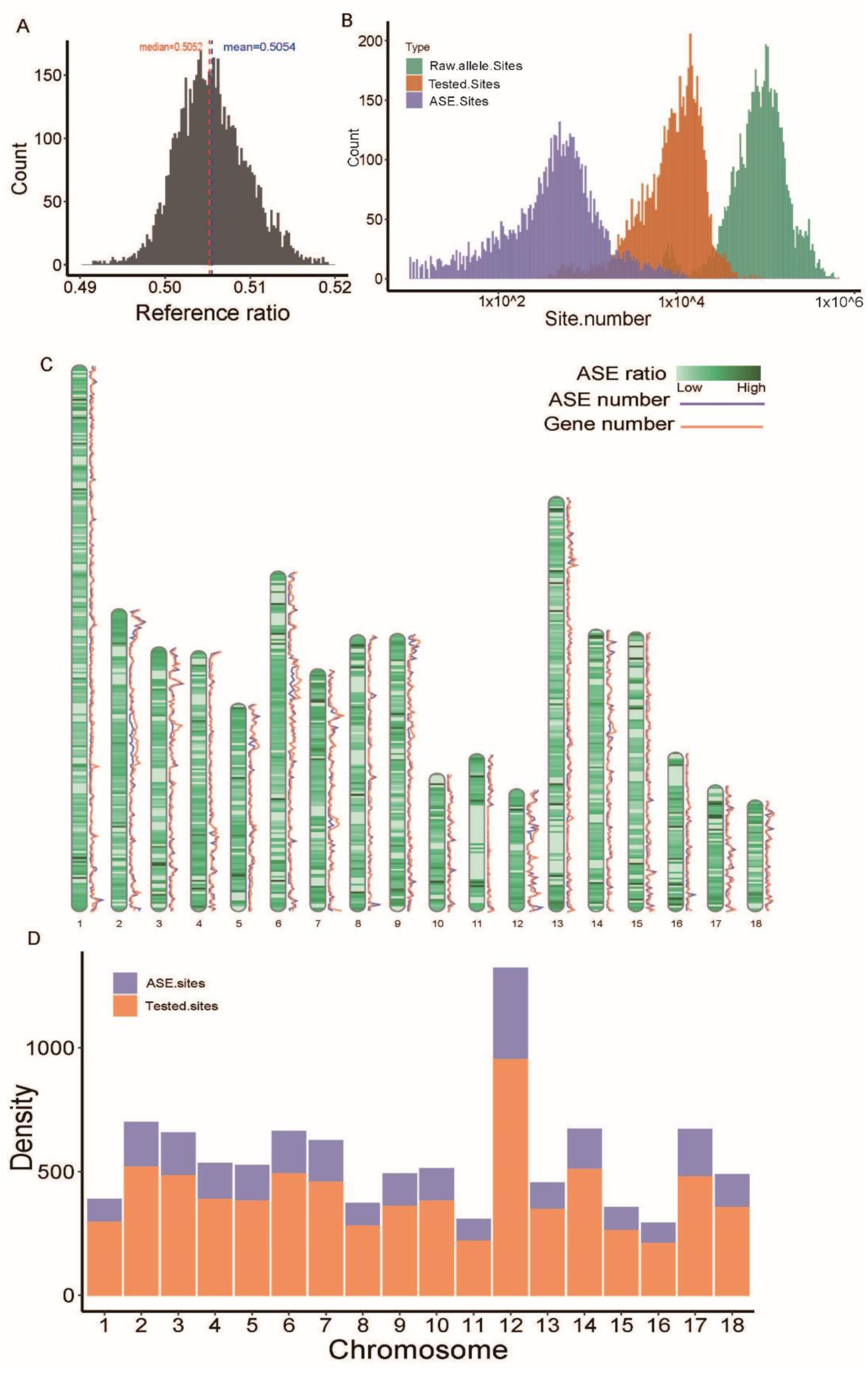
The statistics and distribution of ASE. A). Distribution of the reference ratio for all samples. The mean reference ratio among all samples is shown by the blue dashed line (mean=0.5055), and the median reference ratio for all samples is shown by the red dashed line (median=0.5022). The reference ratio range between 0.491, 0.519. B). The distribution of different allele sites counts among all samples. Raw.Allele.Sites: The raw allele sites number from the phASER software, with at least 1 read covered; Tested.Sites: allele sites passed the applied filtration criteria and were used for the binomial test; ASE sites: the significant ASE sites (FDR <=0.05) after the binomial test and Benjamini-Hochberg corrected. C). The ASE distribution along the genome and the segment’s colors corresponds to the ASE ratio in 1Mb windows (ASE ratio = ASE.Sites / Tested.Sites). The dark purple line indicates the ASE number in 1Mb windows. The orange line indicates the gene number in the 1Mb window. D). ASE site and tested sites density normalized by chromosome length in 1Mb fragments.

Similar patterns were observed when normalized by gene and exon lengths (Additional file 1: Fig S3C, D). To further examine the ASE site occurrences along chromosomes, we captured the accumulated occurrence rate of 1 Mb segments with tested sites and ASE sites for each chromosome (Additional file 1: FigS3E). The accumulation of ASE and tested sites occurred at a similar pace for all autosomes, implying a uniform occurrence of ASE sites along the genome.

For gene-level ASE, a total of 13,684 genes were detected as significant ASE genes with at least 1 ASE site in at least one individual. For each individual, a median of 261 ASE genes was identified among 2,145 tested genes, ranging from 1 to 5,978 ASE genes (Additional file 3: Table S3). The distribution of the ASE genes along the genome showed a similar trend as ASE sites (Additional file 1: Fig S4A). Six segments presented an enrichment of ASE genes, with more than 50 ASE genes. As expected, the distribution of the ASE genes was highly correlated with ASE sites (R^2^=0.85, *p-value* < 2.2e-16, Additional file 1: Fig S4B). We also compared our ASE genes with published imprinted genes in the pig. The majority of published imprinted genes (21 of 30 published imprinted genes) were also found as monoallelic expression genes (MAE gene, Additional file 3: Table S4), which indicated the consistency and reliability of our ASE gene detection. Meanwhile, these imprinted genes were also identified as ASE in certain samples (Additional file 3: Table S4). These results also suggest that these imprinted genes are not strictly mono-allelic expressed.

### Effect of covariates on ASE identification

In RNA-data analyses, the effect of technical covariates can be a considerable problem, especially in gene expression analyses, in which read counts across samples are compared. However, in ASE analyses, read counts from the different alleles at the same position within an individual are compared, which makes it less susceptible to technical covariates [24]. We explored the effect of different technical covariates on the ASE detection. First, we analyzed the correlation between the proportion of significant ASE sites and genes and various technical variates. A relatively high correlation was observed with the number of reads after wasp filtering (adjusted R^2^=0.379), which is expected as the statistical power of detecting ASE is related to the total read count. For the remaining technical covariates, the correlations were significant (*p*-value < 0.05) but not highly correlated (adjusted R^2^ < 0.2). We studied whether the high reads covered on specific sites could minimize the correlation of technical covariates, considering that higher read coverage per site or gene could ensure sufficient detection power. The correlation was decreased with >30 reads covered but remained significant. Also, the correlation difference between the two settings was small (Table 1). To further investigate the effects of technical covariates on ASE patterns across samples, multiple dimensional scale (MDS) clustering of ASE sites was performed based on the pairwise ASE distance between samples. Samples were labeled with different technical covariates. The clustering did not exhibit a specific pattern for the technical covariates across samples (Additional file 1: Fig S5A-C), as contrasted with the tissue and breed clustering (Additional file 1: Fig S5D), which is also consistent with the higher correlation with breeds (adjust R^2^ = 0.292) and tissues (adjust R^2^ = 0.475), indicating the ASE distribution pattern among samples was not driven by batch effect.

**Table 1.** The correlation of ASE with technical covariates.

| | | Reads covered $\geq 15$ | | Reads covered $\geq 30$ | |
| --- | --- | --- | --- | --- | --- |
|  |  | ASE sites | ASE genes | ASE sites | ASE genes |
|  | Model1 | 0.061* | 0.061* | 0.055* | 0.056* |
|  | Platform2 | 0.001 | 0.001 | 0.002 | 0.001 |
|  | Library layout3 | 0.019* | 0.033* | 0.013* | 0.026* |
|  | Mapped reads | 0.184* | 0.251* | 0.090* | 0.194* |
|  | Unique reads | 0.197* | 0.264* | 0.099* | 0.204* |
|  | Wasp filtered reads | 0.379* | 0.410* | 0.247* | 0.340* |
|  | Breed | 0.292* | 0.252* | 0.290* | 0.250* |
|  | Tissue | 0.457* | 0.439* | 0.417* | 0.440* |

\*The adjusted $R^2$ is measured between each covariate and the percentage of significant ASE sites or genes in each sample from GTEx-pig samples with total reads covered over 15 and 30. \* $p$ -value < 0.01. The Spearman correlation was used for continuous covariates, and linear regression for categorical covariates. \*1. Model Contains: BGISEQ-500, Illumina HiSeqXTen, Illumina HiSeq1000(1500, 2000, 2500, 3000, 4000), Illumina NovaSeq 6000, Illumina NextSeq 500, PacBio RS II, ABI SOLID System, Ion Torrent Proton, PacBio RSII. 2. Platform contains: BGISEQ, ILLUMINA, ION\_TORRENT, ABI SOLID, PACBIO SMRT. 3. Library layout: Paired and Single.

### ASE distribution among Tissues and Breeds

Considerable variation was observed in the ASE distribution across tissues and breeds. Among the 42 tissues (3 cell lines included), the ASE percentage ranged from 1.31% in the pituitary to 53.24% in the blastocyst (Fig 2A). The lowest ASE percentage was found in the brain (2.56% ASE on average among brain tissues), consistent with the observation in human GTEx data [11]. Also, the intestine showed a relatively low ASE percentage. A high ASE percentage was found in embryo-related tissues, with the highest in early embryo development. At the gene level, similar trends were observed, albeit with a higher gene ASE percentage than the ASE site (ranging from 4.46% in the pituitary to 66.37% in the blastocyst). The IPEC cell line also exhibited a high ASE percentage at both the SNP and gene levels, which is consistent with the finding that IPEC cell lines have increased ploidy [25] and high-level ASE [26]. At the breed level, breeds with more than 10 samples were selected to explore ASE (cell lines were excluded). As shown in Fig 2B, the percentage of ASE sites and gene fractions greatly varied between breeds. A relatively high ASE percentage was found in the local Chinese breeds (∼18.64% on average). There was no significant difference in the ASE percentage between crossbreeds and commercial breeds (Yorkshire, Landrace, Duroc) (10.2% vs ∼9.05%).

**Fig. 2.**
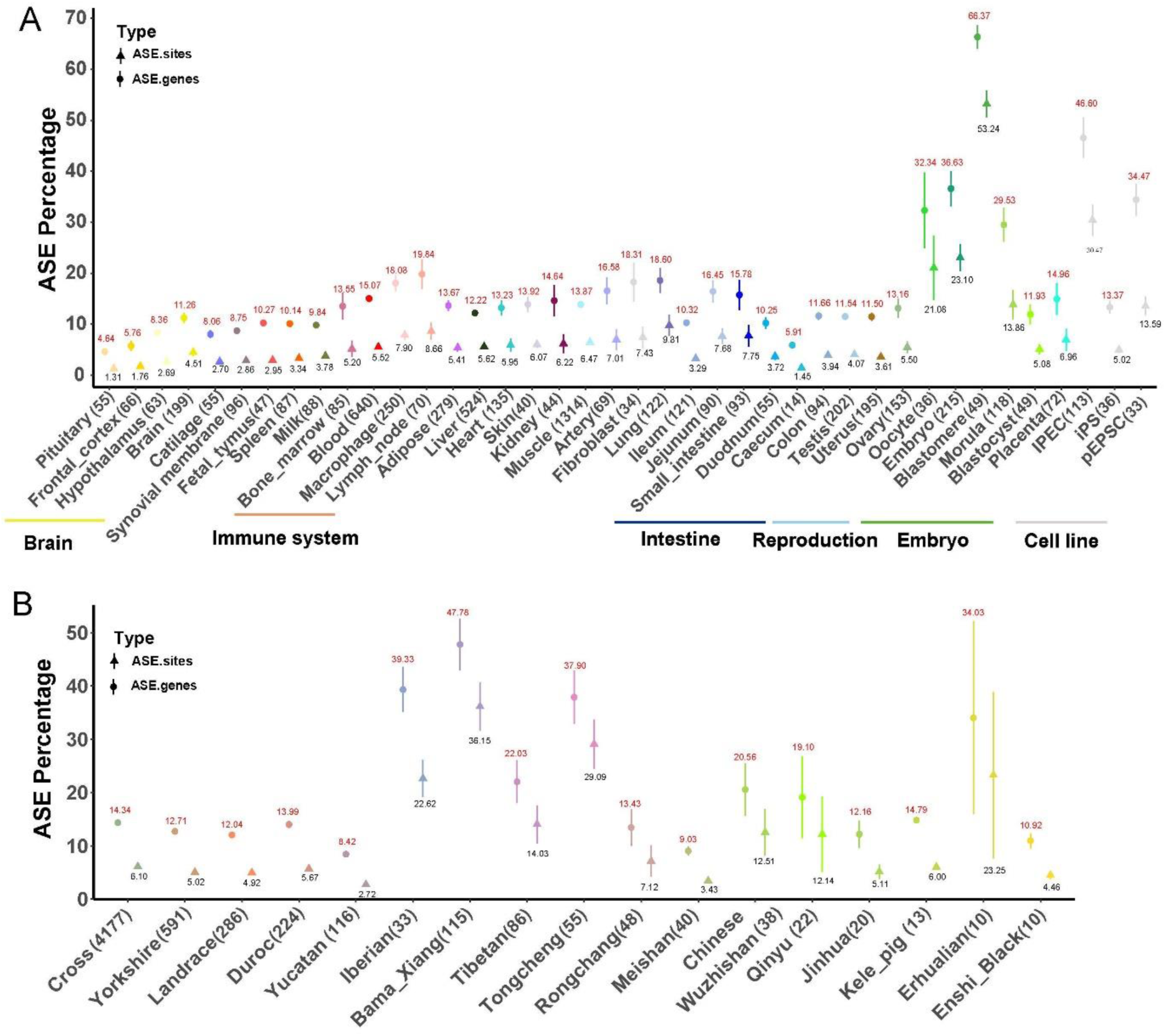
The ASE and ASE gene percentage among tissues and breeds. A). The percentage of ASE in each tissue (mean ± 95% CI). The tissues are labeled in different colors. B). The percentage of ASE in different breeds (mean ± 95% CI). The breeds are labeled in different colors. The circle indicates the ASE gene percentage, with the red number indicating the mean value of the ASE gene percentage in each group, and the triangle indicates the ASE site percentage, with the black number indicating the mean value of the ASE site percentage. The number in the parentheses indicates the sample size.

### Tissue-specific ASE genes

Most ASE genes are expected to result from genomic variation in regulatory elements, and ASE will, therefore, depend on the segregation of these variants within and between populations. In addition, tissue level or tissue-specific ASE, regulated by, e.g., epigenetic marks, may indicate variation in tissue-specific regulatory sites (i.e., enhancers). To explore if tissue-level or tissue-specific ASE genes were present in our data, we searched for genes that were expressed in multiple tissues but only showed ASE in a single or subset of tissues (see material/methods for details). Twenty-eight of the 40 tissues had such ASE genes (Fig 3A & Additional file 3: Table S5). Among these 28 tissues, a total of 165 tissue-level ASE genes were detected (Table S6), ranging from 1 to 63 genes per tissue (7 ASE genes on average). There were 142 tissue-level ASE genes, only detected in one tissue (Fig 3A, B, Additional file 4: Table S6). Among these tissue-level ASE genes, two genes showed strong tissue-specific ASE: ENSSSCG00000011237 (*PDCD6IP*) in Blastomere and ENSSSCG00000017089 (*ANXA6*) in Fibroblast. These were detected as generally tested genes in 28 tissues but only identified as ASE in these two tissues separately. *PDCD6IP* codes for the programmed cell death 6 interacting protein involved in cytokinesis after mitosis. It has been well studied in the mouse, showing ubiquitous expression [27], and involved in multiple processes, such as membrane repair, cytokinesis, and cell proliferation [28]. ANXA6 belongs to a family of calcium-dependent membrane and phospholipid-binding proteins. The gene ENSSSCG00000001229 (*SLA-3*) showed ASE across multiple tissues (in 10 of the 20 tissues). *SLA-3* is an MHC class I gene that plays an indispensable role in controlling adaptive immune responses by recognizing foreign antigens [29].

**Fig. 3.**
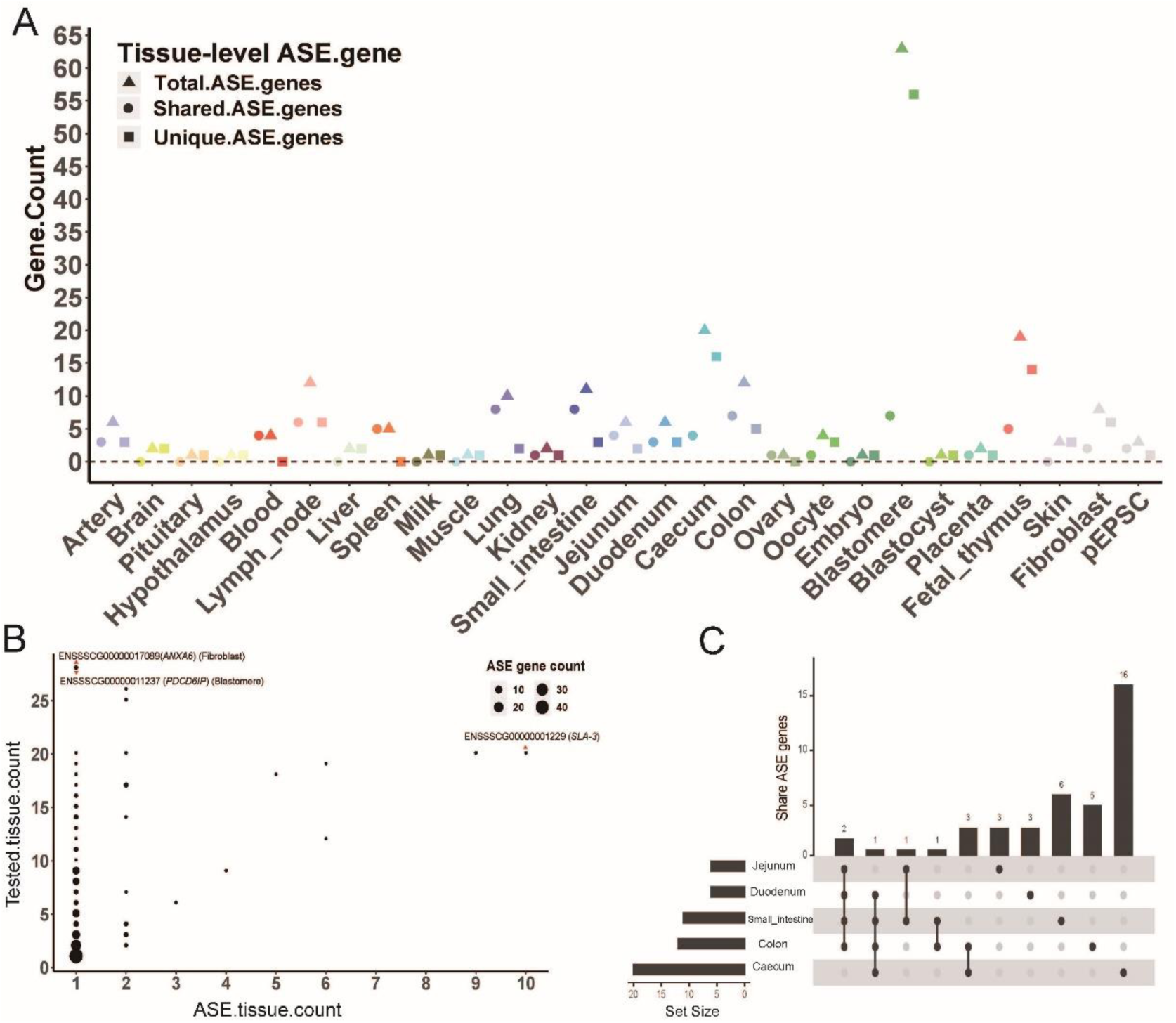
The statistic of tissue-level ASE genes. A). Tissue-level ASE gene number in each tissue. The total ASE genes (triangle) indicate all ASE genes found in each tissue. The shared ASE genes (circle) indicate tissue-level ASE genes detected in more than one tissue. The unique ASE genes indicate that ASE genes are only detected by this tissue. The tissues are labeled in different colors. B). For each ASE gene, the number of tissues that were tested (Y-axis) and the number detected as tissue-level ASE (X-axis). Each point indicates a tissue-level ASE gene. The X-axis represents the number of tissues with this gene as the ASE gene. The y-axis represents the number of tissues that test this gene, and the point size indicates the number of ASE genes tested and detected by a certain number of tissues. The genes ENSSSCG00000017089 (ANXA6) and ENSSSCG00000011237 (PDCD6IP) were detected by 28 tissues and were identified as ASE genes in Fibroblast and Blastomere. The gene ENSSSCG00000001229 (SLA-3) was detected in 20 tissues and detected as an ASE gene in 10 tissues. C). The intersection of tissue-level ASE genes among intestine-related tissues.

We grouped tissues with similar functions to explore the tissue-level ASE signatures further. There were 3 tissue-level ASE genes detected by four intestine-related tissues: *SLA-3*, *OAS2,* ENSSSCG00000041364 (Fig 3C). ENSSSCG00000041364 is a gene with an unknown function. *OAS2* codes for a 2’-5’-oligoadenylate synthetase 2 protein, which is involved in host anti-PRRSV response [30]. As mentioned before, SLA-3 plays a critical role in immune response. These three tissue-level ASE genes were widely expressed in most pig tissues and also detected as ASE genes in other tissues, indicating that these genes are not intestine-specific ASE genes. Across the intestinal tissues, thirty-three tissue-level ASE genes were identified, each detected in at least one tissue but not uniformly across all (Fig 3C). The function for these ASE genes revealed several critical genes involved in different biological processes, such as lysozyme activity (*LYZ, CPSF6*), peptidoglycan muralytic activity (*LYZ, CPSF6*), defence response to bacteria (*LYZ, S100A14, FCN2, CPSF6, OAS1*), vitamin, mineral digestion and absorption (*SLC46A1*) and metabolism (*SUCLG2*). In the brain tissues, four brain-specific ASE genes were identified (*ALDH5A1, GH1, DMP1, DSPP*). *ALDH5A1* is involved in the gamma-aminobutyric acid metabolic process; mutation of this gene is related to neurological phenotypes [31]. GH1 is a member of the somatotropin prolactin family of hormones and is primarily synthesized and secreted in the anterior pituitary [32]. DSPP and DMP-1 are highly phosphorylated proteins and essential for properly developing hard tissues [33].

## Discussion

This study is the first to investigate global ASE across pig tissues and breeds using RNA-seq data compiled by the Farm-GTEx consortium [22]. The large sample size allowed us to establish an extensive ASE resource covering most pig tissues and breeds. We explored the overall distribution of ASE across the genome, tissue-level, and breed-level ASE characteristics.

In this work, an ASE analysis workflow using current best practices was adapted, including WASP filtering to decrease mapping bias and phASER to produce SNP-level and gene-level ASE data. The imputed genotypes created by the pig Farm-GTEx consortium were used to remove mapping bias and perform ASE analysis. First, we confirmed the value of RNA-imputed genotype data by comparing its’ efficiency in mapping bias removal and ASE detection with WGS genotype data. Additionally, we compared published imprinted genes with our ASE results. The consistency of the imprinted genes with our MAE genes confirmed the validity of our analyses. We also observed that some of these known imprinted genes exhibit ASE across various tissues, showing non-strict mono-allelic expression. However, new MAEs cannot be detected for the current study due to an elevated number of genotype errors in the RNA-imputed genotype data, which will result in false MAEs if not filtered out as done in the current study. For the technical covariates, the MDS clustering analysis did not show a significant batch effect caused by these technical covariates. Thus, We used this workflow, applied to thousands of pig samples, yielded relatively widespread ASE resource data using RNA-imputed genotype data. The workflow proves to be suitable for other species where only RNA-seq datasets and imputed genotype datasets are available.

ASE sites were widely but unevenly distributed across the genome, with hotspots correlating with gene-rich regions on the chromosomes. In human LCL cell lines, ASE-rich genomic loci were identified that are particularly associated with immune gene clusters, such as the major histocompatibility complex (MHC) on chromosome 6, where a sharp increase in ASE cumulative occurrence rates was observed [34]. However, in our study, ASE sites exhibited relatively uniform occurrence rates, and no specific functional enrichment was detected within ASE hotspots. This discrepancy may be attributed to the mixed tissue types and the diverse breeds analysed in our study, potentially obscuring the cell-specific patterns observed in more homogeneous datasets.

We explored the diversified ASE distribution across tissues and breeds. Firstly, a distinct clustering pattern of ASE sites was observed on the MDS plot comparing pairwise ASE distance between individuals. Distinct tissue clusters strongly suggest that tissue type has the most considerable effect on the percentage of ASE. Among the tissue-level ASE distribution, the embryo-related tissues showed the highest ASE and different stages of embryo development had variable ASE percentage distribution. Blastomere showed the highest ASE, and during embryo development, the ASE percentage decreased in the morula and blastocyst. It is expected that the paternal genome is silent to avoid rejection by the maternal immune system during the early stages of embryo development [35]. Another high ASE percentage was observed in oocytes. The formation of an oocyte is a complex process involving changes in ploidy, and mature oocytes contain only one allele. Our bulk RNA dataset likely represented multiple oocytes, leading to an elevated ASE percentage. The lowest ASE percentage was found in the brain-related tissues, a similar trend as observed in humans (2% ASE), supposing reduced gene diversity in the brain [36]. However, another study showed that the ASE genes in the porcine brain are estimated to be 42% of all genes [21] and around 14-26% in the cattle brain [2]. Also, in the thymus, the cattle study showed a 10% ASE site percentage (986 ASE sites out of 9,781 tested sites), whereas 2.95% ASE sites was observed in our study. A direct comparison, however, is not appropriate since the differences in the level of ASE may not only come from biological reasons, such as tissue type, species, development stage, genetic diversity, but also from technology confounders, such as different methods, filtering criteria, and statistical ways to detect ASE. Thus, a standard, well-acceptable pipeline could help to reduce the technical bias and improve the comparability of results. The tissue-level ASE profiles tended to be highly individual and tissue-specific, and ASE genes shared within and across tissues and cell types were limited. We still found several tissue-level and tissue-specific ASE genes, and these genes’ functions were consistent with the tissue type. Since epigenetic signals are likely to regulate these ASE genes, future analyses, including haplotype-specific epigenetic data, such as DNA methylation from Nanopore sequencing, may provide clues on how these ASE genes are regulated.

We also explored the breed-level ASE. For some breeds, tissue type effect on ASE percentage partly explained the high ASE percentage, such as for the Bama-Xiang population (47.78%), in which 75 of 115 samples are from blastomere tissue. Besides the tissue effect, population genetic differentiation also has an impact on the ASE sites, as shown by the relatively higher correlation between breeds and ASE percentage (Table 1, adjusted R^2^=0.292). Also, in the human population, the distinct ASE pattern were found between European and African human populations [37]. Chinese local breeds, known for their higher genetic diversity, exhibited relatively high ASE percentages (∼18.64%). In contrast, there was no significant difference in the ASE percentage between crossbreeds and commercial breeds (Yorkshire, Landrace, Duroc) (14.34% vs 12.91%). Several factors may obscure the influence of genetic diversity on ASE levels. For instance, while crossbreeds are expected to have higher genetic diversity and consequently higher ASE percentages, the complex hybrid history between Chinese and Western pigs has shaped the present genomic landscape with elevated ASE percentages in commercial breeds, making them comparable to those in crossbreeds. The introgression from Chinese into European breeds helped to improve economically important traits in commercial breeds [38–40], suggesting that these introgression regions do play a role in gene expression. Thus, future studies applying the origin of alleles (e.g., Asian versus Europa) for ASE genes may provide insight into the effect of historical introgression on gene expression. In our study, we didn’t consider the parent-of-origin effect due to a lack of trio-family samples. Furthermore, the direction of the cross will also affect the ASE expression profile. Studies using reciprocal crosses in chicken [41], mouse [42], and pig [43] have highlighted the complexity of ASE and parent-of-origin expression.

Knowledge of ASE may provide further insights into the molecular basis underlying complex phenotypes. With the vast ASE resource described in the current study, assessing the connection between economically relevant phenotypes and ASE sites/gene profiles could improve the power to find causal variants through GWAS [34].

Loci exhibiting ASE have been associated with production traits, including intramuscular fat content in pig [44] and meat quality in cattle [45]and pig[46]. Several further investigations can be conducted, e.g. combining ASE with eQTL to fine-map cis-regulatory functional variants [47] and exploring introgressed and deleterious regulatory variation [48]. Combining multi-omic data will provide further information about the contribution of epigenetic variation on gene expression variation in a population [49]. Our publicly available SNP-level and gene-level ASE data will provide a powerful resource complementing the available eQTL data established by the Farm-GTEX-pig consortium [22].

### Conclusions

In this work, We used the data from the Pig-GTEx consortium to conduct a vast SNP-haplotype- and gene-level ASE resource consisting of 42 tissues and 34 pig breeds. At first, we observed that the ASE is broad but unevenly distributed among the genome. Then, We explored the diversified ASE distribution across tissues and breeds. The embryo and reproduction-related tissues showed a relatively high ASE percentage. This vast ASE resource has numerous uses for studying regulatory variation, such as studying the effects of rare regulatory variation, fine-mapping cis-regulatory functional variants with the eQTL, and exploring the introgressed regulatory variation. By making the vast SNP- and gene-level ASE data in porcine genome for the first time, we assume that this resource will find similarly broad use as the human’s ASE resource[1, 50].

## Methods

### Sample collection, RNA-seq, and genotype data

In total, 7,008 samples’ RNA sequence data were collected from the porcine Farm-GTEx project [22]. A similar QC workflow was performed as the porcine Farm-GTEx project. Briefly, adaptors were trimmed, and poor-quality reads were discarded using Trimmomatic (v0.39) [51]. Bam files were produced by mapping RNA-seq fastq files to the Sscrofa11.1 reference genome (Ensembl v100) using STAR (v2.7.0) [52], with the options:-outFilterMismatch-Nmax 3,-outSAMmapqUnique 255. Then, the samples with a mapping rate lower than 85% were selected and tested for possible contamination or wrongly assigned species. 10,000 reads were randomly selected from the fastq files from these samples by seqtk (version:1.3-r106) and blasted against the NCBI’s nucleotide (NT) database (NCBI v5) [53]. The samples in which more than 3% of selected reads showed the highest similarity to at least one non-pig species within the top 5 NT-hit species were excluded from the analysis. Reference mapping bias was eliminated by applying the WASP methodology [54] using STAR with options “-waspOutputMode SAMtag” combined with each sample’s genotype data. Finally, 6,927 samples were kept with QC quality ≥ 30 and mapping rates ≥ 65%. The unique (MAPQ=225) and passed WASP filtered (with the tag “vW:i:1”) reads were kept by using the Samtools (Version: 1.14) [55] with the “view” function for the next steps. The GATK (v4.2.1.0) [56] MarkDuplicate mode with default parameters was used to remove PCR duplicate reads.

The imputed, phased genotype data used in this study for removing mapping bias and the ASE analysis, were from the Farm-GTEx consortium and contained 3,087,268 population-level variants from 7,008 samples [22]. In brief, these variants were called from RNA sequence data, and heterozygous SNPs were imputed to genome-wide SNPs based on the multibreed pig genomics reference panel (PGRP, Farm-GTEx). We use the term RNA-imputed genotype for these genotype data. To get sample-level genotype data, Bcftools (Version: 1.9) [55] was used to select the individual genotype data from the RNA-imputed genotype data. The reference homozygous sites (with“0|0”) and clustered SNPs (≥3 SNPs in 10bp windows) were removed to get the final individual genotype. Finally, each sample had an average of 1,400,235 SNPs.

### DNA-seq variant detection

WGS data of 3 individuals from three breeds (Duroc, Pietrain, and Yorkshire) were additionally included (PRJEB19268). The alignment was performed using BWA-MEM (Burrow-Wheeler Aligner) [57] with options “-M-R ‘@RG\tID:PigWUR150\tSM: PigWUR-150-1\tPL:ILLUMINA’”. Freebayes (v0.9.21) [58] was utilized to identify variants for each sample with the options “--min-base-quality 10”, “--min-mapping-quality 20”, “--min-alternate-fraction 0.2”, “--haplotype-length 0”, “--pooled-continuous--ploidy 2”, “--min-alternate-count 2”. Subsequently, VcfFilter (v0.2) [59] was employed to remove low-quality sites using the option “-f QUAL > 20”, while vcfkeepgeno [59] was used to select specific FORMAT fields: “GT DP AD” within the VCF file. Indels were excluded, and bi-allelic SNPs were chosen using VCFtools (0.1.16) [60]. Afterward, clustered SNPs (≥3 SNPs in 10bp windows) were eliminated by GATK (v4.2.1.0).

### Mapping bias filtration and ASE detection

A two-step approach was employed to decrease reference mapping bias. Initially, the option “waspOutputMode” in STAR (v2.7.0) was used, as described above. Despite the use of WASP a preference of outliers towards the reference genome was still observed. The following approach was therefore used to omit these biases: the means of reference ratio was calculated [*Reference ratio = reference reads/total reads*] for each sample. The reference ratio’s Interquartile Range (IQR) across all samples was calculated (IQR=0.0065). Thresholds for selecting non-biased samples were computed as [0.4915, 0.51915] ([Q1-1.5*IQR, Q3+1.5*IQR]). Samples falling outside these thresholds were considered potential outliers and removed for the next analysis.

SNP-level ASE was generated using the phASER (version 1.1.1) package [61] with filtered RNA alignment file and sample genotype file described above as input file with the following settings:-mapq 255 and -baseq 10. The raw allele sites were obtained from the phASER output for each sample. Subsequently, the sites were selected by these filter criteria for further analysis: 1) total mapped reads ≥ 15 for each locus/sample; 2) minimum of 3 reads mapped to reference or alternative allele; 3) proportion of mapped reference or alternative reads accounting for > 2% of the total mapped reads. These filtering criteria removed the sites with potential sequencing errors and mono-allelic expression (MAE). MAE sites were excluded due to the inflated number of MAE sites caused by errors in imputed genotype calling (unpublished observation). For the remaining sites (termed as ‘tested sites’ hereafter), a binomial test was used to estimate significant deviation from the expected 0.5 allele ratio (with the binom.test() function in R) [62]. Raw *p* values were corrected for multiple tests via the Benjamini-Hochberg test [63]. The ASE sites with a false discovery rate (FDR) ≤ 0.05 were considered significant.

Haplotype-level ASE was generated using the phASER package. For each sample, the haplotype expression output file (haplotypic_counts.txt) contains the number of unique reads that map to each haplotype for all phased haplotype blocks. Then, haplotypic blocks with a total read count of more than 8 were selected for the ASE gene analysis. The gene-level data were computed using phASER Gene AE 1.2.0, employing--min_haplo_maf 0.01, and utilizing the Ensembl annotation Sscrofa11.1 v104 gene features bed file as well as the filtered haplotypic count file from phASER. The raw output of the ASE gene contained the unique read count and variant number for each gene per sample. For the raw gene-level ASE data, similar processes were used to select the ASE gene as for ASE sites: 1) the total number of allelic reads covering each gene is ≥ 15 reads; 2) the allelic reads mapped to each haplotype are ≥ 2; 3) the proportion of reads for each haplotype accounts for > 2% of the total mapped read. The remaining genes (termed ‘tested gene’) were tested for significant imbalanced expression using binomial tests, and *p* values were corrected using the Benjamini-Hochberg method, and ASE genes with FDR ≤ 0.05 were considered significantly imbalanced expressed.

Lastly, only samples with more than 200 tested sites and 10 ASE sites were retained for further analysis, resulting in 6,022 samples (**Table S1**). The workflow of the ASE analysis pipeline is detailed in FigS1(Additional file1: Fig.S1).

### ASE distribution

To investigate the distribution of SNP level and gene-level ASE along the pig genome, the chromosomes were segmented into 1 Mb windows, and ASE site numbers and ASE ratio (ASE ratio = ASE sites/tested sites per 1 Mb) were calculated accordingly. The ASE number on each chromosome is related to the chromosome length. To further investigate the ASE enrichment difference on each chromosome, the ASE density per 1Mb was calculated per chromosome and normalized based on the length of each chromosome, total gene length and total exon length. The ASE density is the normalized number of ASE site in a 1 Mb segment, which was calculated as follows:

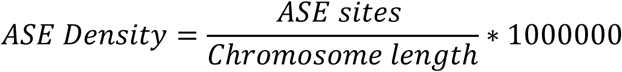

### Multidimensional scaling cluster by ASE data

The classical multidimensional scaling (MDS) was used to evaluate sample clustering. At first, the distance between the two samples was calculated as the proportion of non-shared ASE sites at the total shared tested sites [24]. Then, a pairwise distance matrix was built for all samples, and MDS was produced by the cmdscale() function in R (v4.2.3).

### General tissue-level ASE gene identification

Gene ASE was calculated at the sample level. To find ASE genes that are general ASE in a tissue or subset of tissues, we performed the following steps. Firstly, the small sample size tissues with similar functional anatomy were merged into one category, such as cerebellum (n=2) and cerebrum (n=2), which were merged into the brain tissue category. Then, for each tissue category, the following criteria were applied: 1) A tested gene (see definition above) is detected in at least 70% of samples in a specific tissue category (i.e., the gene is generally expressed in the tissue), and a sample with this tested gene is defined as a tested sample; 2) at least 70% of the tested samples showed significant ASE for the gene. Genes fulfilling these two criteria are termed tissue-level ASE genes and if only detected as ASE in one tissue, these genes are termed tissue-specific ASE genes. Cell lines were excluded from this analysis. GO analysis (ShinyGO 0.80 [64]) was performed for all tissue-level ASE genes per tissue to explore these ASE gene functions.

## Declarations

### Ethics approval and consent to participate

Not applicable

### Consent for publication

Not applicable

### Availability of data and materials

All RNA raw data analyzed in this study are publicly available for download without restriction from SRA (https://www.ncbi.nlm.nih.gov/sra/) and BIGD (https://bigd.big.ac.cn/bioproject/) databases. Details of RNA-seq can be found in Supplementary Table1. All the computational scripts used for RNA-seq alignment, quantification, quality control, and ASE analysis are available at the FarmGTEx GitHub website (https://github.com/FarmGTEx/PigGTEx-Pipeline-v0), and The imputed genotype file can be accessed from Farm-GTEx, PigGTEx-Portal (https://piggtex.farmgtex.org/). **The raw ASE datasets generated during this study for each sample are available**

### Competing interests

The authors declare that they have no competing interests

## Funding

This work was supported by grants from the Shenzhen Outstanding Talents Training Fund Grand No. 202102, National Key Research and Development Program of China (Grant No. 2021Y FD301201, 2024YFF0728800), National Natural Science Foundation of China (Grant NO. 31972539, 31501933), STI 2030—Major Projects (Grant NO. 2022ZD04017), National Natural Science Foundation of Guangxi province (Grant NO.2024GXNSFAAO10105), Science, Technology, and Innovation Commission of Shenzhen Municipality (Grant NO. JCYJ20180306173644635), Science Technology Innovation and Industrial Development of Shenzhen Dapeng New District (Grant NO.PT20170201, PT202101-05).

## Authors’ Contribution

L.K., L.B. and L.F. conceived and designed the study. WY.Y. conducted the analyses and wrote the manuscript. L.K., L.B. contributed to the data and computational resources. M.G., O.M., M.A.M.G., and L.F. provided the valuable suggestions for the analysis. L.K., L.B., M.G., O.M., M.A.M.G., and L.F. revised and edited the manuscript. All authors discussed the results, read and approved the final paper.

## Acknowledgments

We would like to thank all groups of the Farm-GTEx-Pig Consortium for their support of this project. We thank the GTEx members for their valuable suggestions and contributions to the data collection.

## Supplementary information

**Additional file 1**: Supplemental Figures S1-S5. (docx 1798KB)

Fig S1: The workflow of Allele-Specific Expression analysis applied to the Farm-GTEx-pig dataset.

Fig S2: A) The impact of removing mapping bias using different genotype data; B) The number of the ASE sites detected by different genotype data, C),D),E): The final sample size and reference ratio among different tissues and breeds

FigS3: The statistics of ASE sites.

FigS4: A). The distribution of ASE gene and ASE sites across the genome; B). The correlation between ASE genes and ASE sites.

Fig S5: Multidimensional scaling (MDS) clustering of all ASE samples

**Additional file 2**: Supplemental Table S1-S2(xlsx. 477KB)

Table S1:All Samples(6022) information for ASE analysis.

Table S2: The comparison of the SNPs and ASE site counts by different genotype data.

**Additional file 3** Supplemental Table S3-S5 (docx,83KB)

Table S3: Basic Statistics of ASE Analysis

Table S4: Published imprinted gene

Table S5: Tested gene and ASE gene number at tissue level

**Additional file 4** TableS6 tissues-levels ASE gene (xlsx, 18KB)

## Notes

### Competing Interest Statement

The authors have declared no competing interest.

