## supplemental FigS1-S5, TableS3-S5 for "Mapping the Landscape of Allele-Specific Expression In Porcine Genomes"

### Supplemental Figures

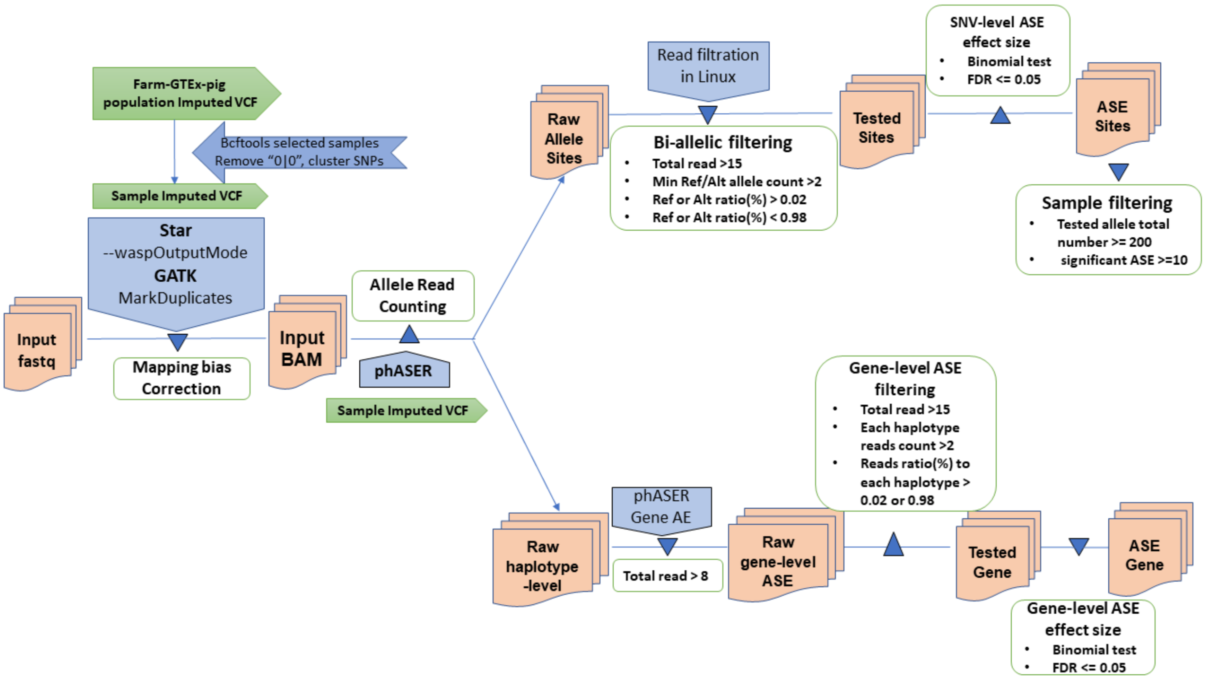


**Fig S1**: **The workflow of Allele-Specific Expression analysis applied to the Farm-GTEx-pig dataset.** The imputed, phased VCF files from the GTEx consortium were used to filter out reference mapping bias and conduct ASE analysis. The mapping and reference mapping bias filtration were performed using STAR (with --waspOutputMode), and GATK (Genome Analysis Toolkit, v4.2) was utilized to identify duplicates. The phASER tool packet was employed for SNV-level and gene-level ASE analysis. A binomial test was conducted to assess the significant imbalance in the expression between two allelic sites (or haplotypes).

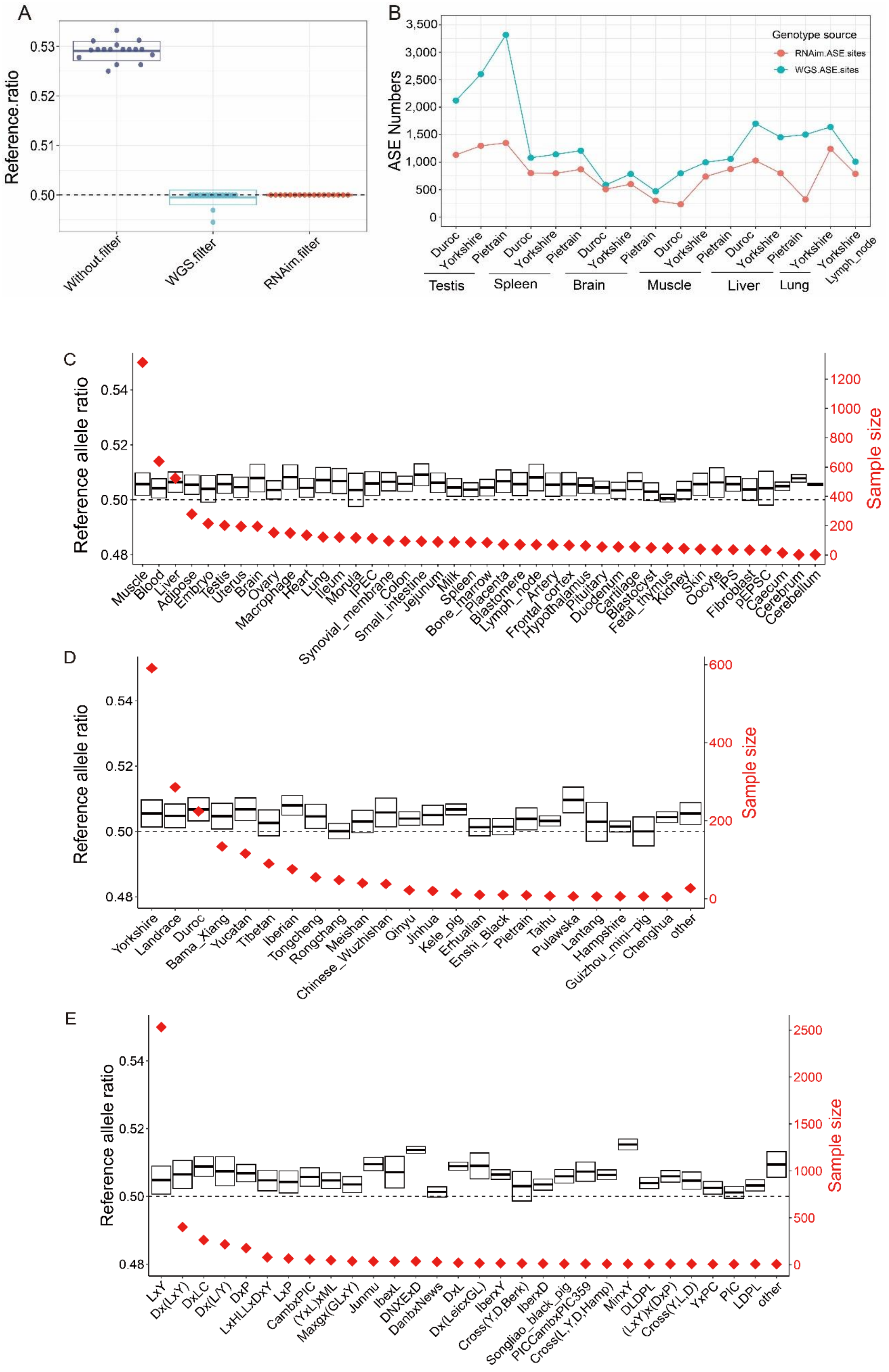
A
Reference.ratio
0.53
0.52
0.51
0.50
Without.filter
WGS.filter
RNAim.filter
B
ASE Numbers
3,500
3,000
2,500
2,000
1,500
1,000
500
0
Genotype source
RNAim.ASE.sites
WGS.ASE.sites
Duroc
Yorkshire
Pietrain
Duroc
Yorkshire
Pietrain
Duroc
Yorkshire
Pietrain
Duroc
Yorkshire
Pietrain
Duroc
Yorkshire
Pietrain
Yorkshire
Yorkshire
Lymph_node
Testis
Spleen
Brain
Muscle
Liver
Lung
C
Reference allele ratio
0.54
0.52
0.50
0.48
Sample size
1200
1000
800
600
400
200
0
Muscle
Blood
Liver
Adipose
Embryo
Testis
Uterus
Brain
Ovary
Macrophage
Heart
Lung
Ileum
Morula
IPEC
Synovial_membrane
Colon
Small_intestine
Jejunum
Milk
Spleen
Bone_marrow
Placenta
Blastomere
Lymph_node
Artery
Frontal_cortex
Hypothalamus
Pituitary
Duodenum
Cartilage
Blastocyst
Fetal_thymus
Kidney
Skin
Oocyte
iPS
Fibroblast
pEPSC
Caecum
Cerebrum
Cerebellum
D
Reference allele ratio
0.54
0.52
0.50
0.48
Sample size
600
400
200
0
Yorkshire
Landrace
Duroc
Bama_Xiang
Yucatan
Tibetan
Iberian
Tongcheng
Rongchang
Meishan
Chinese_Wuzhishan
Qinyu
Jinhua
Kele_pig
Erhualian
Enshi_Black
Pietrain
Taihu
Pulawska
Lantang
Hampshire
Guizhou_mini-pig
Chenghua
other
E
Reference allele ratio
0.54
0.52
0.50
0.48
Sample size
2500
2000
1500
1000
500
0
LxY
Dx(LxY)
DxLC
Dx(L/Y)
DxP
LxHLLxDxY
LxP
CambxPIC
(YxL)xML
Maxgx(GLxY)
Junmu
IbexL
DNXExD
DanbxNews
DxL
Dx(LeicxGL)
IberxY
Cross(Y,D,Berk)
IberxD
Songliao_black_pig
PICCambxPIC359
Cross(L,Y,D,Hamp)
MinxY
DLDPL
(LxY)x(DxP)
Cross(Y,L,D)
YxPC
PIC
LDPL
other

**Fig S2**: A). The impact of removing mapping bias using different genotype data (without.filter = STAR align only without correction, WGS.filter = genotype data is from WGS sequence, RNAim.filter = genotype data is from imputed RNA sequence). B). The number of the ASE sites detected by different genotype data. C),D),E): The final sample size and reference ratio among different tissues C), pure breeds D), and crossbreeds E) after filtering out mapping bias. The red diamond represents the sample size of the target group. The boxplot shows the reference ratio for each group. In C), the "other" group consisted of breeds with a sample size of less than 5 (including Dapulian: 4, Alentejano: 4, Yanan: 3, Wannanhua: 3, Laiwu: 3, Anqing six-end-white: 3, Qingping: 2, Jiaxing_black_pig: 2, Diannan_Small_Ear: 2). In E), the Cross.other group contained the groups with a sample size of less than 5 (including PxY: 3, ExS: 2, Lx(G/D): 2). The full name of each group can be found in TableS1.

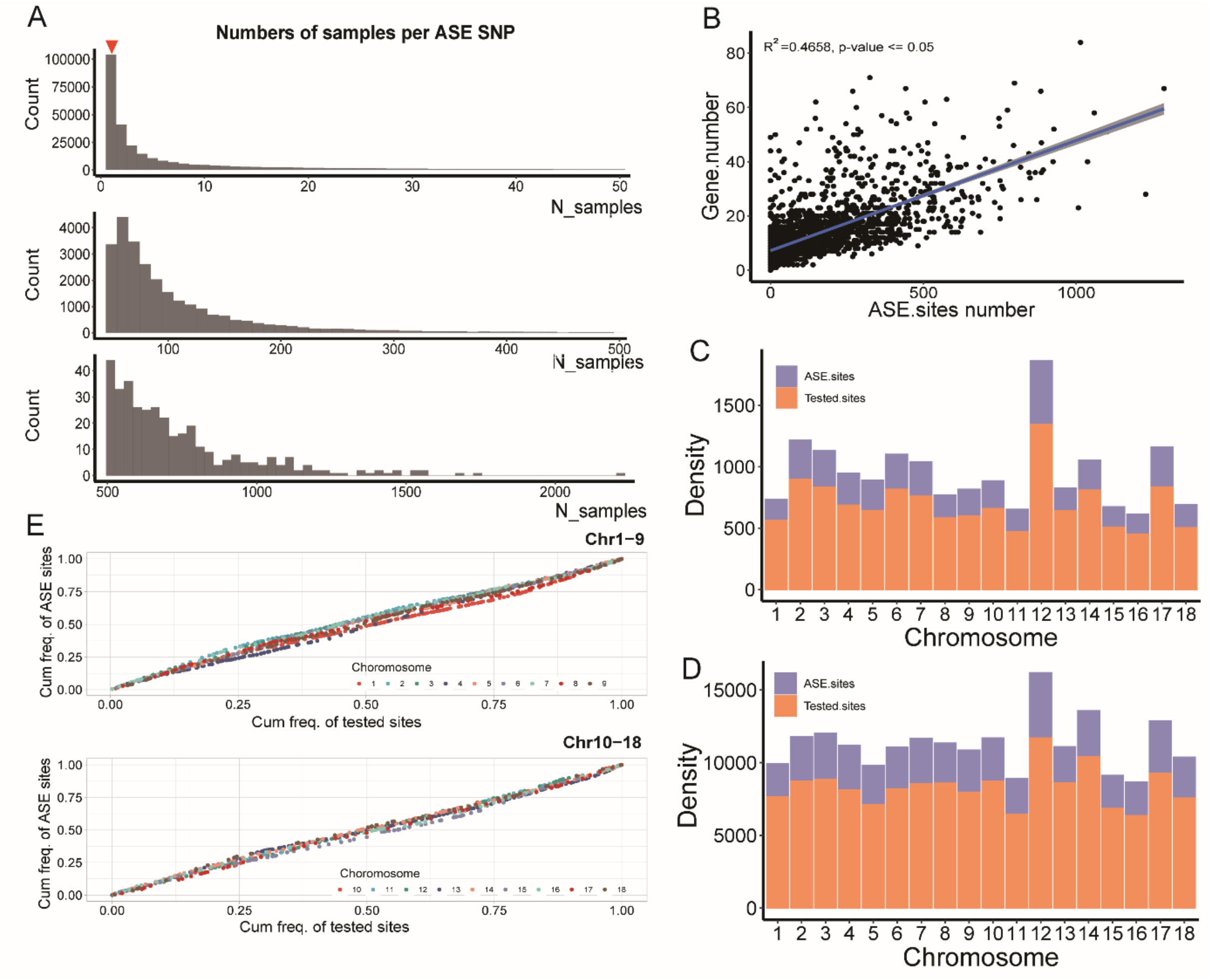


**Fig S3**: **The statistic of ASE sites** A) Histogram of the number of samples for the ASE SNPs. The red triangle means there are 104,078 ASE sites only detected in one sample. B) The correlation between the gene number and significant ASE sites in the genome. C) D). The normalized ASE sites and tested sites density per 1Mb among the chromosomes. C) Normalized by gene length in each chromosome; D) Normalized by exon length in each chromosome. E) The cumulative fraction of ASE sites and tested sites per 1Mb segments in each chromosome.

FigureS4

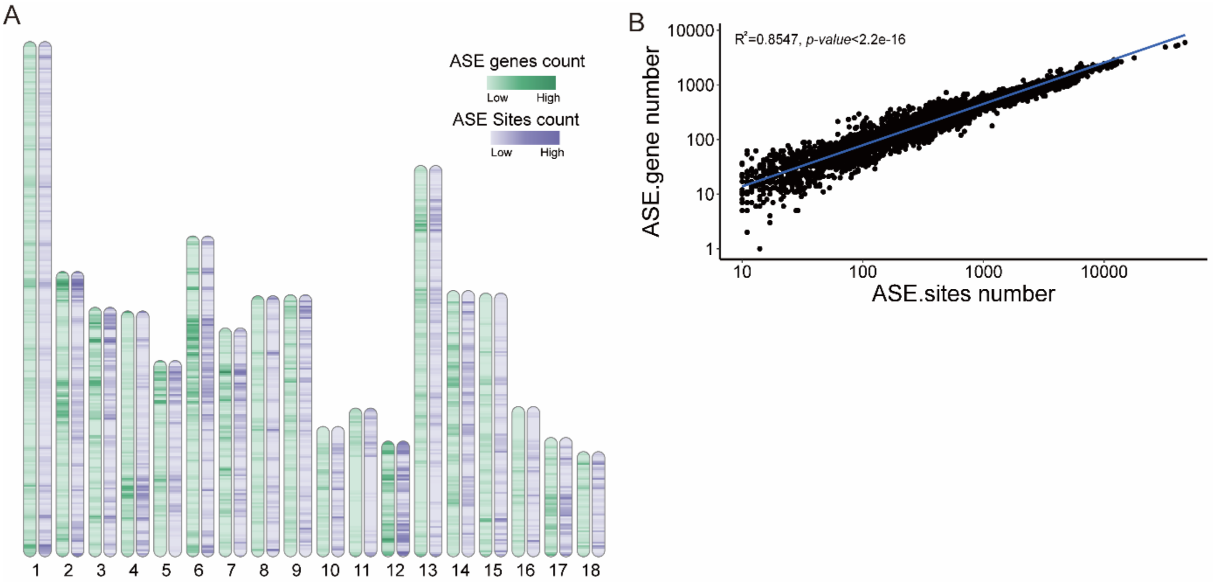


**Figure S4**: A). The distribution of ASE gene and ASE sites across the genome. The segments' colors correspond to the number in 1Mb windows. Green: ASE genes, purple: ASE.Sites. B). The correlation between ASE genes and ASE sites. The adjusted R-squared is 0.8547; the p-value is < 2.2e-16.

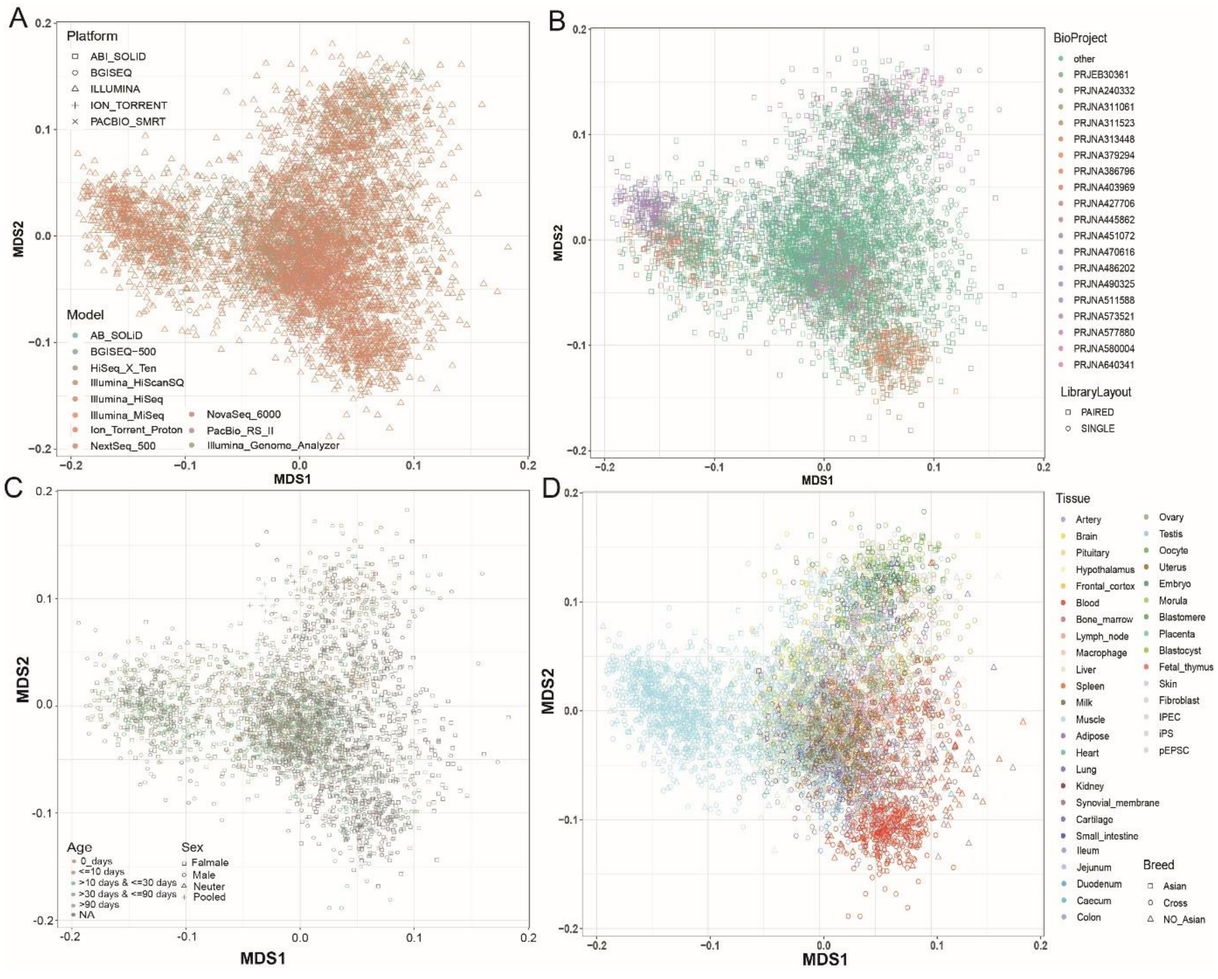


**Figure S5: A-D** Multidimensional scaling (MDS) clustering of all ASE samples. Samples are colored and labeled by different technical covariates (as shown in Table 1). A) Samples are labeled by sequence platform and colored by different sequence machine models. B). Samples are labeled by the Library layout (single or paired reads) and colored by different Bioproject ID. There are 393 Bioprojects involved, and we only colored the bioprojects with more than 50 samples(19 Bioprojects). The remaining samples were grouped into the “other” group, which is green in the plot. C) Samples were labeled by sex and colored by age. For each age group, we categorized the samples into 6 groups according to the days after birth (age = 0 days; age ≤ 10 days; age > 10 days & ≤ 30 days; age > 30 days & ≤ 90 days; age > 90 days and NA). D) Samples are colored by tissues and labeled by breeds. Breeds were categorized into Asian, non-Asian, and Cross groups (Table S1).

### Supplemental TableS3-S5

##### **Table S3**: Basic Statistics of ASE Analysis

| | Tested sites (≥ 15 reads) | ASE sites (FDR < 0.05, ≥ 15 reads) | ASE fraction* (%) | Tested genes (≥ 15 reads) | ASE genes (FDR < 0.05, ≥ 15 reads) | ASE gene fraction* (%) |
|---|---|---|---|---|---|---|
| Minimum | 268 | 10 | 0.23 | 81 | 1 | 0.42 |
| Mean | 11,316 | 850 | 7.49 | 2,144 | 342 | 15.87 |
| Median | 10,091 | 458 | 4.55 | 2,143 | 261 | 11.93 |
| Maximun | 92,052 | 46,825 | 77.22 | 7,750 | 5,978 | 88.12 |

*ASE fraction=(ASE sites/ Tested sites)*100

##### **Table S4**: Published imprinted gene

| | Gene symbol | Ensemble ID | ASE samples count (FDR <=0.05) | MAE gene samples count | Tested sample count | reference |
|---|---|---|---|---|---|---|
| 1 | *DIRAS3* | ENSSSCG00000003797 | NA | NA | NA | [1, 2] |
| 2 | *DIS3L2* | ENSSSCG00000022177 | 49 | 190 | 1366 | [3] |
| 3 | *DLK1* | ENSSSCG00000035805 | NA | NA | NA | [4, 5] |
| 4 | *FABP3* | ENSSSCG00000036883 | 293 | 60 | 1378 | [6] |
| 5 | *GATM* | ENSSSCG00000004672 | 117 | 28 | 370 | [7, 8] |
| 6 | *GNAS* | ENSSSCG00000007520 | 387 | 204 | 1518 | [3, 9] |
| 7 | *IGF2* | ENSSSCG00000035293 | NA | NA | NA | [5, 6] |
| 8 | *IGF2R* | ENSSSCG00000004044 | 885 | 328 | 1600 | [3, 5] |
| 9 | *JPH3* | ENSSSCG00000002653 | 39 | 34 | 253 | [3] |
| 10 | *KBTBD6* | ENSSSCG00000031378 | 129 | 65 | 1001 | [3] |
| 11 | *KCNQ1* | ENSSSCG00000039556 | 65 | 98 | 413 | [3] |
| 12 | *LTA* | ENSSSCG00000001403 | NA | 1 | 23 | [6] |
| 13 | *MC5R* | ENSSSCG00000028411 | NA | 1 | 9 | [6] |
| 14 | *MEST* | ENSSSCG00000016554 | 54 | 107 | 221 | [5] |
| 15 | *NAP1L5* | ENSSSCG00000009209 | NA | NA | NA | [5, 10] |
| 16 | *NNAT* | ENSSSCG00000007336 | 86 | 38 | 354 | [1, 5, 11] |
| 17 | *NOB1* | ENSSSCG00000002753 | 496 | 131 | 3415 | [3] |
| 18 | *PEG10* | ENSSSCG00000036049 | 3 | 8 | 15 | [5, 8] |
| 19 | *PEG3* | ENSSSCG00000003326 | 4 | 8 | 17 | [5, 10, 12] |
| 20 | *PGM2L1* | ENSSSCG00000014838 | 466 | 71 | 1919 | [3] |
| 21 | *PHLDA2* | ENSSSCG00000021597 | 36 | 3 | 86 | [5] |
| 22 | *PLAGL1* | ENSSSCG00000004127 | 45 | 55 | 134 | [5] |
| 23 | *RTL1* | ENSSSCG00000002573 | NA | NA | NA | [4] |
| 24 | *SGCE* | ENSSSCG00000015328 | 10 | 10 | 29 | [5] |
| 25 | *SGIP1* | ENSSSCG00000025243 | NA | NA | NA | [3] |
| 26 | *TACC2* | ENSSSCG00000037530 | 328 | 31 | 2150 | [3] |
| 27 | *THRB* | ENSSSCG00000036033 | 128 | 15 | 1383 | [3] |
| 28 | *UOX* | ENSSSCG00000022206 | 167 | 58 | 1712 | [6] |
| 29 | *WT1* | ENSSSCG00000013316 | NA | NA | NA | [6] |
| 30 | *ZNF709* | ENSSSCG00000013715 | 31 | 24 | 287 | [3] |

**Table S5**: Tested gene and ASE gene number at tissue level.

1. The tested gene number is the number of tested genes in each tissue. It is counted when the gene is detected as the tested gene (see Methods) by at least 70% of samples in this tissue, and the sample with this tested gene is called the tested sample.

*2 The ASE gene number is the number of the ASE genes in each tissue. It is counted when the gene is detected as an ASE gene by at least 70% of the tested samples.

*3 Unique ASE gene number: the number of unique ASE genes which was only detected in this tissue.

| Tissue | Tested gene number[1] | ASE gene number[2] | Unique ASE gene number[3] |
|---|---|---|---|
| Artery | 129 | 6 | 3 |
| Brain | 212 | 2 | 2 |
| Pituitary | 66 | 1 | 1 |
| Hypothalamus | 443 | 1 | 1 |
| Frontal_cortex | 182 | 0 | 0 |
| Blood | 90 | 4 | 0 |
| Bone_marrow | 97 | 0 | 0 |
| Lymph_node | 411 | 12 | 6 |
| Macrophage | 65 | 0 | 0 |
| Liver | 54 | 2 | 2 |
| Spleen | 350 | 5 | 0 |
| Milk | 140 | 1 | 1 |
| Muscle | 80 | 1 | 1 |
| Adipose | 53 | 0 | 0 |
| Heart | 168 | 0 | 0 |
| Lung | 559 | 10 | 2 |
| Kidney | 189 | 2 | 1 |
| Synovial_membrane | 8 | 0 | 0 |
| Cartilage | 13 | 0 | 0 |
| Small_intestine | 853 | 11 | 3 |
| Ileum | 20 | 0 | 0 |
| Jejunum | 334 | 6 | 2 |
| Duodenum | 270 | 6 | 3 |
| Caecum | 880 | 20 | 16 |
| Colon | 422 | 12 | 5 |
| Testis | 37 | 0 | 0 |
| Uterus | 133 | 0 | 0 |
| Ovary | 185 | 1 | 0 |
| Oocyte | 55 | 4 | 3 |
| Embryo | 25 | 1 | 1 |
| Blastomere | 82 | 63 | 56 |
| Morula | 16 | 0 | 0 |
| Blastocyst | 159 | 1 | 1 |
| Placenta | 285 | 2 | 1 |

| | | | |
|---|---|---|---|
| Fetal_thymus | 1087 | 19 | 14 |
| Skin | 338 | 3 | 3 |
| Fibroblast | 213 | 8 | 6 |
| IPEC | 351 | 136 | 121 |
| iPS | 521 | 10 | 7 |
| pEPSC | 126 | 3 | 1 |
